# The origin, invasion history and resistance architecture of *Anopheles stephensi* in Africa

**DOI:** 10.1101/2025.03.24.644828

**Authors:** Tristan P.W. Dennis, Jihad Eltaher Sulieman, Mujahid Abdin, Temesgen Ashine, Yehenew Asmamaw, Adane Eyasu, Eba A. Simma, Endalew Zemene, Nigatu Negash, Abena Kochora, Muluken Assefa, Hamza Sami Elzack, Alemayehu Dagne, Biniam Lukas, Mikiyas Gebremichael Bulto, Ahmadali Enayati, Fatemeh Nikpoor, Ashwaq M. Al-Nazawi, Mohammed H. Al-Zahrani, Bouh Abdi Khaireh, Samatar Kayed, Abdoul-Ilah Ahmed Abdi, Richard Allan, Faisal Ashraf, Patricia Pignatelli, Marion Morris, Sanjay C. Nagi, Eric R. Lucas, Anastasia Hernandez-Koutoucheva, Patricia Doumbe-Belisse, Adrienne Epstein, Rebecca Brown, Anne L. Wilson, Alison M. Reynolds, Ellie Sherrard-Smith, Delenasaw Yewhalaw, Endalamaw Gadisa, Elfatih Malik, Hmooda Toto Kafy, Martin J. Donnelly, David Weetman

## Abstract

The invasion of Africa by the Asian urban malaria vector, *Anopheles stephensi,* endangers 126 million people across a rapidly urbanising continent where malaria is primarily a rural disease. Control of *An. stephensi* requires greater understanding of its origin, invasion dynamics, and mechanisms of widespread resistance to vector control insecticides. We present a genomic surveillance study of 551 *An. stephensi* sampled across the invasive and native ranges in Africa and Asia. Our findings support a hypothesis that an initial invasion from Asia to Djibouti seeded separate incursions to Sudan, Ethiopia, and Yemen before spreading inland, aided by favourable temperature, vegetation cover, and human transit conditions. Insecticide resistance in invasive *An. stephensi* is conferred by detoxification genes introduced from Asia. These findings, and a companion genomic data catalogue, will form the foundation of an evidence base for surveillance and management strategies for *An. stephensi*.

---

*Anopheles stephensi* is a primary malaria vector native to South Asia and the Persian Gulf(*1*). Well adapted to urban settings, it was discovered in Djibouti in 2012, and has since been discovered in the wider Horn of Africa (HoA), Yemen, and Kenya, in an apparently ongoing range expansion. *An. stephensi* is a highly competent vector of *Plasmodium falciparum* and *Plasmodium vivax* malaria, and has been associated with substantial increases in malaria cases in Djibouti(*2*) and Ethiopia(*3*). The potential spread of *An. stephensi* to cities across Africa - to an estimated 126 million malaria-naïve people(*4*) - has raised international alarm(*5–7*). This has spurred urgent efforts to understand and contain its expansion, particularly in the HoA and Yemen - a region facing geopolitical instability, millions of displaced and vulnerable people, and high susceptibility to the effects of climate change(*8*). Identifying the origin of *An. stephensi*, and the landscape and environmental factors facilitating its spread would enable targeted vector control and surveillance at points of entry, and the development and refinement of models predicting both the spread of the mosquito and of potential implementation of biological and gene drive control techniques. Furthermore, identifying threats to insecticide-based control methods via genomic surveillance data will enable more focused and proactive resistance management strategies, supporting more optimal programmes to control and eradicate *An. stephensi* in its invasive range.

Previous genetic analyses of *An. stephensi* have found greater similarity between Asian and African mitochondrial barcodes(*9*, *10*), and a stepping stone pattern of diversity and population in structure moving inward into Ethiopia from the coast(*11*). The relatively coarse resolution of mitochondrial barcodes compared to genomic data(*12*), a focus on specific geographic regions(*11*), often with small sample sizes(*13*), leaves many unanswered questions about the fine-scale structure and invasion history of *An. stephensi* in Africa. Analysing whole-genome-sequence (WGS) data of individuals sampled systematically from the native and invasive ranges provides clarity on the often complex dynamics of biological invasions, such as invasion timing, origin, and the number of introductions(*14*). Genomic surveillance studies are transforming the evidence base for the control of both invasive species(*14*) and malaria vectors, based on the analysis of vector population structures, the distribution of insecticide resistance mechanisms and gene drive targets(*15–20*). Here, we present a genomic surveillance study of 551 *An. stephensi* specimens sampled from 9 countries across the invasive and native ranges (**Figure 1a, Table S1, Supporting Material**). We present evidence of the likely origin and routes of the *An. stephensi* invasion, characterise the genomic basis of resistance, and provide a foundational genomic data resource for use in combating the challenge to malaria control presented by *An. stephensi*.

**Figure 1.**
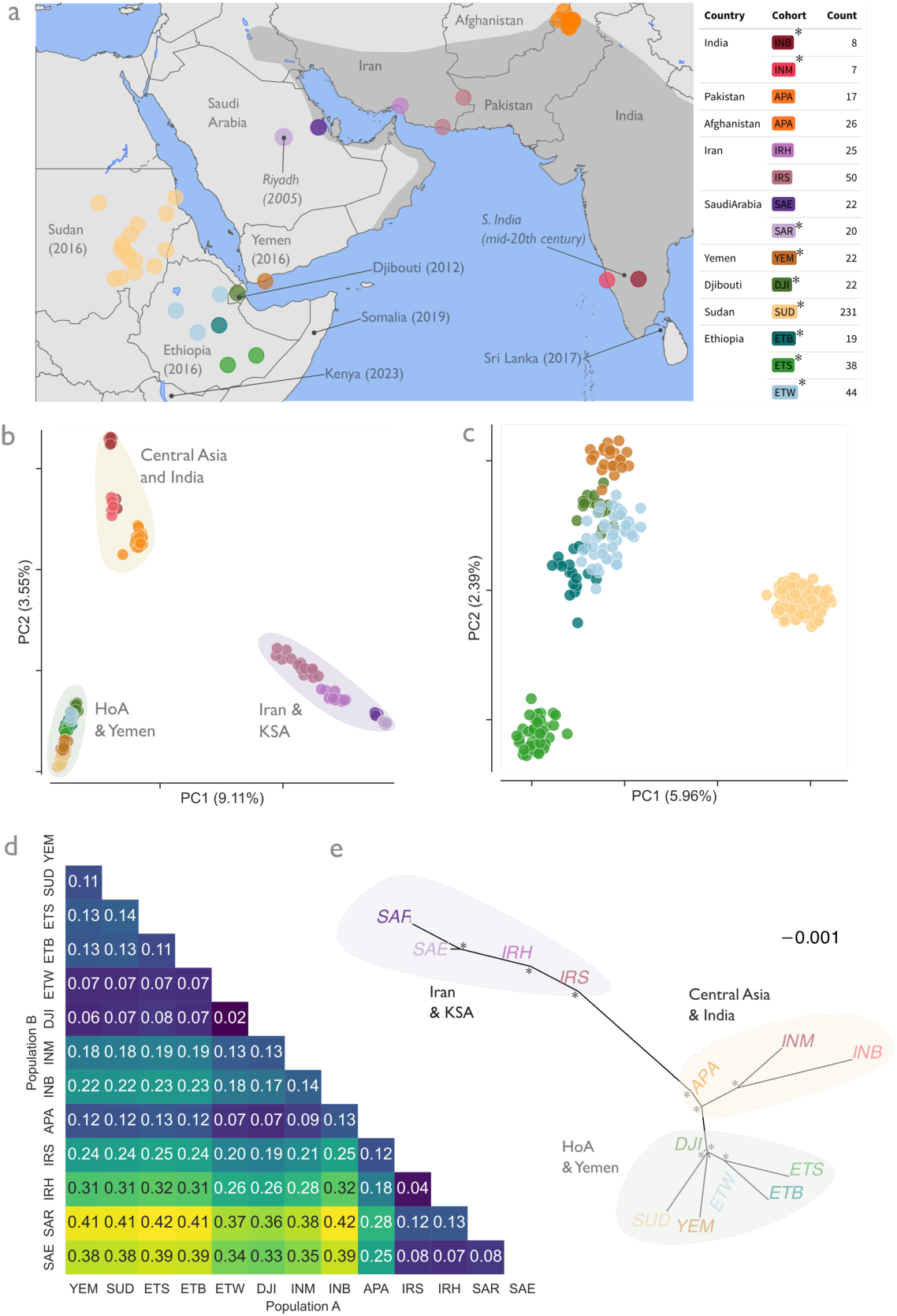
Global population structure suggests an Asian origin of *An. stephensi*. **(A)** map showing sequenced individuals grouped into 12 analysis cohorts based on collection location, principal component analysis (PCA) (**B**) and neighbour-joining tree (**Figure S2**) clustering. Shaded region denotes the native range(*1*), with date of detection in invaded countries and regions (italics). PCA of global samples (**B**) was downsampled to include no more than 30 individuals per cohort, with shaded regions indicating super-population. **(C)** a PCA of samples from the HoA and Yemen only. In both PCAs, points are coloured by cohort, and were generated using 1e5 randomly selected SNPs from accessible regions of chromosome 2. (**D**) Heatmap of mean Hudson’s *Fst* calculated between cohorts, where 0 indicates no differentiation, and 1 indicates complete differentiation. *Fst* was generated using SNPs from accessible regions of chromosome 2. (**E**) A consensus unrooted admixture graph, produced by resampling 1e5 accessible SNPs 1000 times from chromosome 2. Shaded regions denote super-populations, with tips labelled and coloured by cohort. Node labels indicate bootstrap support, and scale bar indicates drift parameter.

## An Asian origin of African An. stephensi

*An. stephensi* has undergone repeated global range expansions: migrating from Northern to Southern India in the mid-20th century(*21*), westwards across the Arabian Peninsula in the early 2000s, and from an unknown origin to Africa in 2012 (**Figure 1a**). Since first detection in the Port of Djibouti(*2*), *An. stephensi* has been detected across the HoA region (**Figure 1a**)(*5*, *22–26*), with possible transient reported occurrences in Nigeria (2020)(*5*) and Ghana (2023)(*27*). WGS of 551 *An. stephensi,* including collections across the HoA and Yemen between 2021-2023, (**see Supporting Material and Table S1 for more detail on collection methods, dates, and locations**), indicate that *An. stephensi* is genetically structured into three population groups across its range: Iran and Saudi Arabia (KSA), Central Asia and India, and the HoA and Yemen (**Figure 1b, Figure S1**). Variation in genetic structure is dominated by the split between the KSA cohorts, and those from Central Asia and Africa. This signal of population structuring is consistent across the *An. stephensi* autosomal genome (**Figure S3),** with no sign of variation characteristic of large polymorphic inversions, in contrast to *An. funestus* and *An. gambiae*(*15, 28*). Afghanistan and Pakistan are the closest related population to those in the HoA and Yemen across both the autosomal chromosomes (**Figure 1d)**, and the mitochondrion (**Figure S4**). Shared selective sweeps and profiles of copy number variation at metabolic insecticide resistance gene families further support a close relationship between Afghanistan and Pakistan and HoA and Yemeni *An. stephensi* (**Figure 2a, b, Figure S5, Table S2**). Without further sampling from Pakistan and Northern India, confident inference of the original invasive origin, single or multiple, is challenging, and it is likely that genetic drift brought on by the population bottleneck of the invasion (**Figure 3**) makes HoA and Yemeni *An. stephensi* look even more genetically distinct **(Figure 1b)**. Nevertheless, the relative genetic distance between KSA and the HoA and Yemen **(Figure 1b, d, e, Figure S1, S2, S3**), clearly precludes an introduction from the Arabian Peninsula(*2*, *29*)

**Figure 2:**
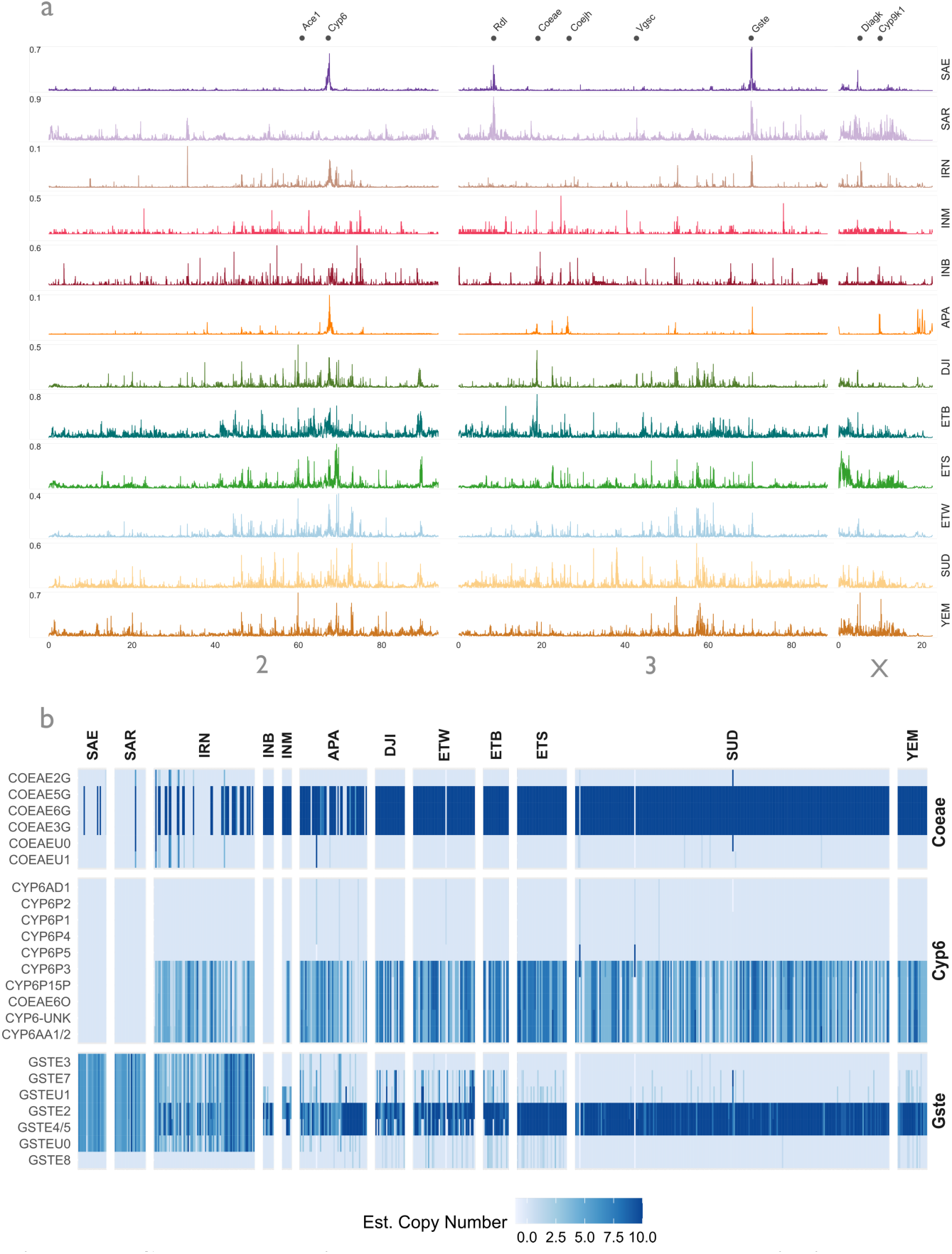
Shared selective sweeps and copy number variation suggest a metabolic basis of resistance, shared between African and Asian *An. stephensi*. **A)** genome-wide selection scans (GWSS) using Garud’s H12 statistic. Peaks indicate localised regions of reduced haplotype diversity, which can indicate the presence of a selective sweep. Y axis indicates H12, X axis indicates chromosomal position in 10s of MB. GWSS were performed on phased haplotypes in windows of 5kb. Plots are panelled columnwise by chromosome (label at bottom, see annotations at top for location of resistance genes), and row-wise by cohort (see label). The noisy signal in the HoA and Yemen is a result of numerous ROH (**Figure 2b**). **Table S2** details the presence and sharing of selective sweeps. (**B**) heatmap of estimated copy number variation (CNVs) at selected metabolic resistance loci. Gene families were selected according to the presence of a selective sweep in (**a**), and prior involvement in resistance in other *Anopheles* taxa(*17*, *19*, *58*). CNVs were estimated using a HMM of deviations from modal coverage in read alignment files(*19*). Heatmap columns indicate individual samples, panelled vertically by cohort. Rows indicate genes found in swept regions (**Table S2**), labelled according to the corresponding ortholog in the *An. gambiae* PEST genome(*59*). Genes without an identifiable *An. gambiae* ortholog are labelled with a ‘U’. Cell darkness indicates increasing copy number (see key). **Figure S5** shows further evidence of shared sweeps between populations.

**Figure 3.**
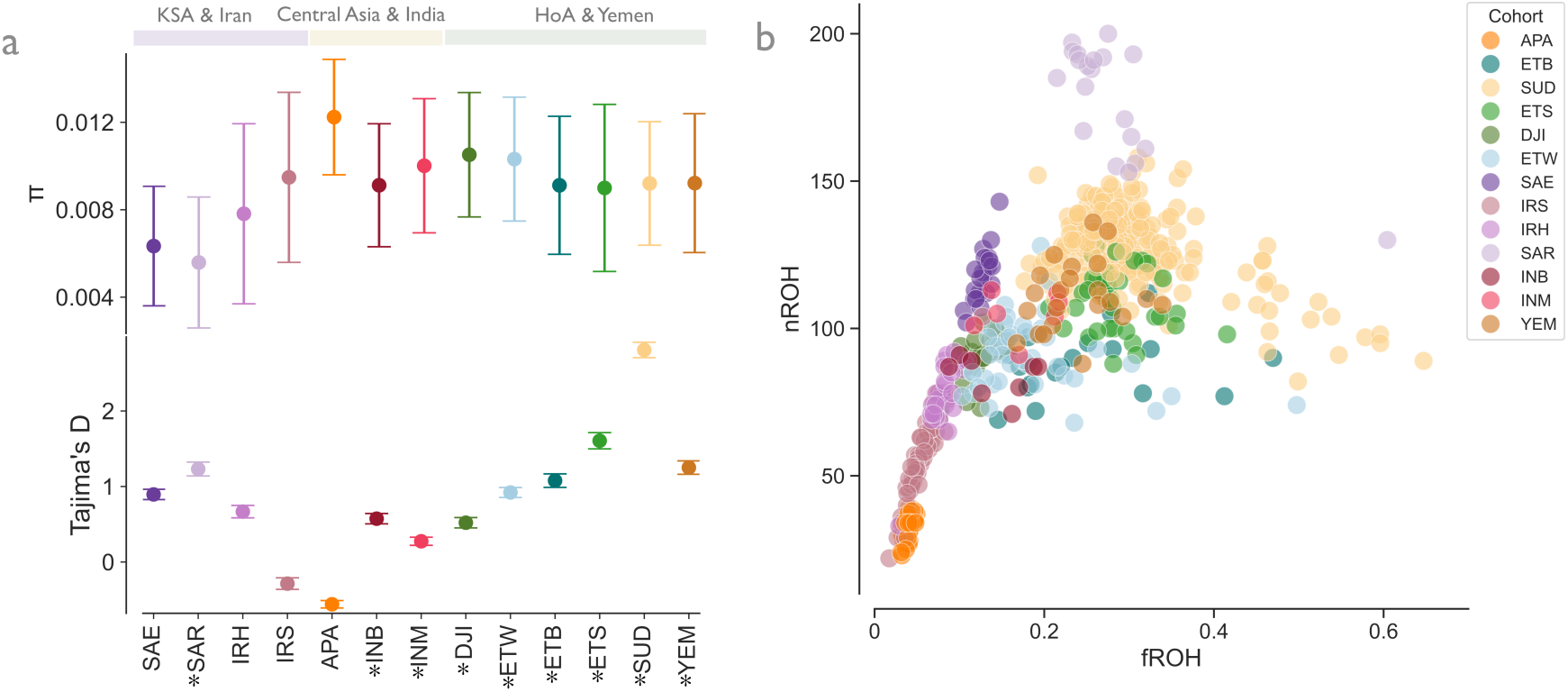
*An. stephensi* has undergone recurrent bottlenecks and diversity loss in multiple invasions. (**A**) shows the nucleotide diversity (π) and (**B**) Tajima’s D of *An. stephensi* calculated using accessible SNPs chromosome 2-wide, with error bars indicating 95% confidence intervals estimated by block-jackknife resampling. Point colour and X-axis label indicate cohort, with invasive populations denoted by asterisks. Shaded bars at top show super-population. (**B**) number (y axis) and fraction of an individual genome (x axis) in runs of homozygosity (ROH). ROH are informative for recent inbreeding, with greater nROH indicating small populations or admixture, and greater fROH indicating recent inbreeding(*34*). ROH were inferred from accessible SNPs genome-wide with a Hidden Markov Model(*15*, *60*), and filtered to include ROH greater than 1e5 in length(*15*, *60*). Points are coloured by cohort (key).

## Spread from Djibouti to the HoA and Yemen

Within each population group, signals of genetic structure and diversity provide an insight into the population history of *An. stephensi* (**Figure 1c, d, e**). Samples from the HoA and Yemen form single groups for Sudan, Southeastern Ethiopia, and Yemen, and a diffuse group containing samples from Djibouti and two populations from West/Central Ethiopia (ETW and ETB) (**Figure 1c, Figure S1)**. These cohorts are tightly clustered together on the global PCA (**Figure 1b**) and neighbour joining trees (**Figure S2**), show low levels of differentiation with respect to one another (**Figure 1d**), form a monophyletic group in an admixture graph (**Figure 1e**), and contain multiple mitochondrial lineages (**Figure S4**). Furthermore, they share selective sweep and CNV profiles at the same metabolic resistance gene families (**Figure 2, Figure S5**), and are derived, to varying extents, from the same ancestry components (**Figure S6**).

Biological invasions may be accompanied by genetic signals of population bottlenecks: reductions in genetic diversity and increased inbreeding(*30*). *An. stephensi* shows signs of extensive diversity loss in multiple populations across the sampled range (**Figure 3a**). The HoA and Yemen, Djibouti and Central Ethiopian cohorts are the most diverse and least inbred. Sudan, Yemen, and Southern Ethiopia are the least diverse, with many individuals from these populations showing signs of intense inbreeding (**Figure 3b**), suggesting that these cohorts represent more severely bottlenecked, or recently established, *An. stephensi* populations.

We infer high levels of migration between Djibouti and West/Central Ethiopia (ETW and ETB), Sudan and Yemen, within Sudan and Djibouti and West/Central Ethiopia (**Figure 4**). Barriers to migration are apparent between Ethiopia and Sudan, Djibouti and West/Central Ethiopia and Southeastern Ethiopia, and Saudi Arabia and the HoA and Yemen (**Figure 4a**). When considering the spatial signal, the closer relationships between Ethiopia, Sudan, and Yemen populations to Djibouti, than to each other (**Figure 1c, d, e, Figure S1**), we aver that the *An. stephensi* invasion of the HoA and Yemen consists of at least three spatially distinct incursions, derived from the initial Djibouti population. Gene flow within the Ethiopian incursion (consisting of samples from Ethiopia and Djibouti City) and Sudanese incursion are significantly higher than gene flow between incursions (**Figure 4b, Table S3, Supporting Methods)**. For the Sudanese and Ethiopian incursions, the further a sample is located from the locations of first detection for these incursions: Port Sudan for Sudan, and Port of Djibouti, for Djibouti and Ethiopia), the less diverse, and more inbred it is likely to be (**Table S3, Figure 4c, d**).

**Figure 4:**
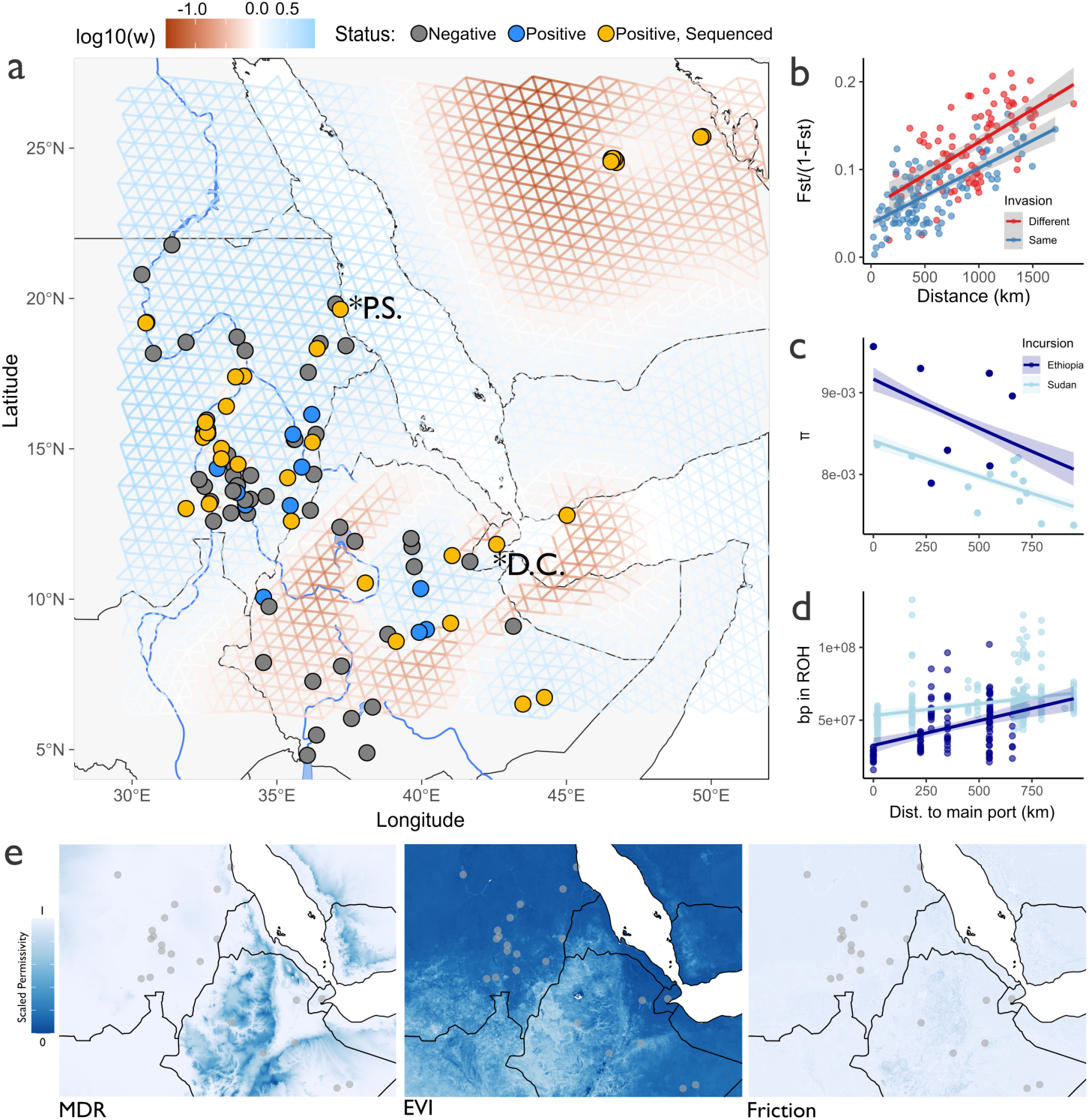
An initial introduction to Djibouti seeded incursions of Yemen, Ethiopia and Sudan in a stepping-stone pattern. (**A**) The grid shows inferred migration (log10(w) across the HoA, Yemen, and Arabian Peninsula. Grid is coloured from dark brown (low) to bright blue (high) between sequenced sampling locations (yellow points). Blue and grey points show where *An. stephensi* was sampled(*61*), and found, or not found respectively. The grid was inferred with fEEMS(*62*), using 1e5 randomly selected SNPs from across Chr2, and selected using leave-one-out cross-validation. Note that edge effects in the model lead to spurious inferences where no data exist (e.g. over Southeastern Ethiopia). Based on the hypothesis (see main text) that initial introduction in Djibouti spawned three geographically separate invasions: into Ethiopia, Sudan, and Yemen, (**B**) shows a pattern of isolation-by-distance in *An. stephensi* in Sudan and Ethiopia/Djibouti, with the y axis showing observed genetic distance (in linearised *Fst*), and the x axis geographic distance, between sampling locations within and between (point colour) invasion subpopulations. (**C**) and (**D**) show how nucleotide diversity (π) and inbreeding (base-pairs in ROH) decrease and increase, respectively, in sampling locations, as distance increases from the main port in Sudan (Port Sudan, “P.S.” on the map) and Ethiopia and Djibouti (Djibouti City, “D.C.” on the map). For **B**, **C** and **D,** solid lines and shading show predicted regression lines and 95% confidence intervals from Gaussian (**A, D**) and gamma (**C**) GLMs. Colours correspond to different levels of interaction variables (see keys). (**E**) show rasters of environmental predictors facilitating *An. stephensi* dispersal: mosquito development rate - MDR, enhanced vegetation index - EVI and transit friction, selected by a Bayesian Gaussian GLM (see main text and methods). Dark - white gradient indicates increasing terrain permissivity (see key).

Previous studies have suggested that human transport links may facilitate *An. stephensi* dispersal in Ethiopia(*11*), and various bioclimatic variables have been linked to *An. stephensi* abundance(*4*, *31*, *32*). We investigated the extent to which eight environmental variables may impact *An. stephensi* gene flow between sampling locations (see **Supporting Material, Table S4**). We applied circuit theory(*33*) to estimate the mean value of selected predictors between sample locations, and fitted general linearised models to test for associations between these environmental covariates and matched linearised *Fst*. These models show that presence of increased enhanced vegetation index (EVI) (mean −.027, 95% CI −.036:-0.019) mosquito development rate (mdr), (mean −.007, 90% CI: −.017:0.001), and decreased landscape travel friction (mean 0.030 90% CI: 0.018:0.044) (**Figure S8**) is associated with *An. stephensi* genetic connectivity (reduced *Fst*). More variation could be attributed to these variables than isolation-by-distance alone (**Figure 4e, Table S5, Supporting Material**), suggesting these factors are important determinants of mosquito spread into the HoA.

## Genetic diversity and structure in the native range

Genetic structure and diversity of *An. stephensi* reflect repeated biological invasions worldwide. Southern Indian *An. stephensi* are less diverse and more inbred than those in Afghanistan and Pakistan (**Figure 3a, b**), reflecting the encroachment of *An. stephensi* southward in the mid-20th century(*21*). Curiously, KSA *An. stephensi* are particularly genetically depauperate (**Figure 3a**). The invasive KSA-Riyadh cohort is less diverse still than the native range in KSA-Eastern (**Figure 3a**), but both KSA cohorts lack the high fraction of the genome in runs of homozygosity (fROH) that would indicate severe inbreeding(*34*) (**Figure 3b**); instead low diversity may be a result of older bottlenecks and population expansion, or relatively mild inbreeding over a long period of time. Despite the relative geographic proximity of Iran to Afghanistan and Pakistan, the two Iranian cohorts are more closely related to KSA than to Central Asia and India, or Africa. The Sistan and Baluchistan cohort (IRS), which is geographically and genetically closest to Afghanistan and Pakistan (APA) (**Figure 1b, d, e, Figure S1, S2, S6**), is the most diverse. The Bandar Abbas cohort (IRH) is much less diverse, more inbred (**Figure 3a, b**), and more closely related to the KSA cohorts (**Figure 1b, 1d, 1e**). CNV and selective sweep profiles in Iran contain gene families found both in Afghanistan and Pakistan, and Saudi Arabia (**Figure 2a,b, Figure S7**): with swept Cytochrome p450 subfamily 6 (*Cyp6*) in IRS shared between Afghanistan, Pakistan, and HoA, and sweeps at the (Glutathione-S-Transferase) *Gste* cluster and *Rdl* being shared between Iran and KSA (**Figure S7**). Low diversity in KSA *An. stephensi* may be a result of relative isolation of the population: the historic range covers a slender region of the east coast of KSA(*1*), <u>where it</u> <u>was first described in 1957</u>(*35*). In a surveillance program beginning in 1971, *An. stephensi* was not detected in Kuwait - between KSA and Iran - until 1985(*36*), raising the possibility that *An. stephensi* in KSA may have been a historical introduction from Iran across the Persian Gulf.

## Evolution and mechanisms of insecticide resistance

Compounding the threat of urban malaria transmission in Africa is widespread resistance of *An. stephensi* to widely used classes of insecticides(*37–40*) - the mainstay of malaria vector control efforts. However, there is limited understanding of resistance mechanisms in the native range(*37*, *41–43*), and especially the invasive range(*13*, *39*, *44*). We screened the genomic data generated in this study for resistance-determinants via a combination of: searching for target-site resistance mutations in known resistance genes (**Figure S7**), performing genome-wide selection scans (**Figure 2a**) and haplotype clustering (**Figure S5)** to identify regions of the *An. stephensi* genome under selection, putatively as a result of insecticide selection pressures. Strong selective sweeps at detoxification enzyme families (**Figure 2a, S5, Table S2**), a lack of selection at target-site resistance genes, and low frequencies of mutations therein (**Figure 2a, S7, Table S2**), combined with extensive copy number variation at metabolic gene families (**Figure 2b**), suggests that resistance in *An. stephensi* in the HoA and Yemen is predominantly mediated through metabolic mechanisms (**Supporting text 1**). Asynchronous selective sweeps at different metabolic resistance haplotypes of the same gene families: *Gste* and *Cyp6,* in KSA, Asia and Africa, suggest convergent evolution through copy number variation at different members of these families may drive resistance in different populations in the absence of gene flow between the Arabian Peninsula and Africa/Asia (**Figure 2, Supporting Text 1, Table S2, Figure S5**).

## On the origin and spread of *An. stephensi* in Africa

Our data suggest that *An. stephensi* in the HoA and Yemen may be derived from an initial bridgehead population in Djibouti. This inference is based on three findings: i) that the population now spreading in Africa is very genetically similar (**Figure 1c, S1**); the presence of Djibouti at the base of a monophyletic clade (**Figure 1e**) of all HoA and Yemen cohorts, and that all are more closely related to Djibouti and to each other, than to any external population (**Figure 1b, c, d, e**); ii) a stepping stone pattern of diversity loss in inland, more recently established, populations. (**Figure 2**, **Figure 4c, d**); and iii) uniform resistance profiles (**Figure 2, Figure S5**). A maritime origin hypothesis(*45*) is supported by the pattern of genetic structuring (**Figure 4a**), and loss of diversity away from Port Sudan and Djibouti (**Figure 4b,c**), as well as first discoveries of *An. stephensi* in coastal cities: Djibouti(*2*), Port Sudan(*5*), Aden City(*24*), Hodeidah(*46*) and Bosaso(*47*). Based on the relative similarity of eastern Iranian, Afghan, Pakistan and HoA/Yemen resistance haplotypes and CNV profiles (**Figure 2b, Figure S5**) a possible scenario is that African *An. stephensi* originated in southern Pakistan. Widespread resistance in the invasive and native range is likely largely mediated by extensive copy number variation at metabolic mechanisms, supported by gene flow and convergent evolution.

To date, hypotheses for the dynamics and structure of the *An. stephensi* invasion have been based largely on phylogenetic and haplotype-network based analyses of short cytochrome c-oxidase subunit I (*COI*) sequences. *COI* phylogenies, and interpretations of high diversity (relative numbers of *COI* haplotypes) are used to propose hypotheses of multiple introductions(*48*, *49*), ongoing connectedness with the native range(*9*) and wind-borne migration of *An. stephensi* as a primary method of introduction and spread(*29*). Our analysis of the *COI* phylogeny failed to resolve any population genetic structure beyond the split between *An. stephensi* from the KSA and Iran, and Central Asia and Africa (**Figure S4b**). Moreover, our evidence of severe genetic bottlenecks in Africa and the Arabian Peninsula (**Figure 2**) contradicts inference of substantial genetic diversity based on the presence of multiple *COI* haplotypes in the invasive range(*9*). Network analysis of reduced-representation sequencing data has been used to propose the eastern Ethiopian city of Dire Dawa as a hub of *An. stephensi* dispersal(*11*). An alternative explanation is spatial autocorrelation in the genetic data due to the central location of Dire Dawa in Ethiopia(*50*). Whilst we did not have samples from Dire Dawa in our dataset, similarity of genetic diversity (**Figure 3a**) and admixture profiles (**Figure S6**) between Djibouti and West/Central Ethiopia samples, and generally high gene flow in the region (**Figure 1d, 4a**) make it difficult to pinpoint a specific hub location. More confident inferences of possible dispersal hubs will require sampling schemes explicitly designed to detect spatial genetic structure(*51*), or specific hypothesised dispersal mechanisms or corridors (e.g. in *Aedes spp.* mosquitoes(*52–55*)). Multi-model comparison and selection can then be used to evaluate the relative fits of models of landscape features or dispersal corridors(*33*, *56*) against alternative models, including isolation by distance.

## A foundational data resource for genomic surveillance

Genomic data are transforming our understanding of the ecology, evolution, and resistance architecture of malaria vectors(*15*, *16*, *19*, *28*). These studies are facilitated by genomic data that are accessible, well-documented, and free to use. In an effort to assist the community with the design of resistance allele surveillance, development of phenotyping assays, and gene drive targets, as well as for comparison with new population genomic data, we have made available a mature catalog of SNP, haplotype, and CNV data in partnership with the MalariaGEN Vector Observatory(*57*). This body of data and analysis will underpin future efforts to predict the spread of *An. stephensi*, manage and monitor resistance, surveil entry points in targeted vector control efforts, and facilitate the control and elimination of invasive *An. stephensi*.

## Acknowledgements

We thank all the sampling teams, local authorities, community leaders and residents involved in specimen collection, the LSTM scientific computing department, as well as Hilary Ranson and Charles Wondji for helpful discussion of the manuscript.

## Author contributions

Conceptualization: DW, MJD, AMR, ALW, HTK, EM, EG, DY

Study design: DW, MJD, TPWD, AMR, RB, AE^2^, HTK, EM, YA, AE^1^, EAS, EG, DY, ALW

Resources: JES, MA, TA, YA, AE^1^, EAS, EZ, NN, AK, MA, HSE, AD, BL, AE^2^, FN, AA, MA, BAK, SK, AAA, RA, PDB

Sample processing: FA, PP, MM, TA, YA, JES, AE^1^, AD

Analysis: TPWD, ESS

Software: ERL, SCN

Data management: TPWD, AH-K

Funding acquisition: MJD, ALW, DW, EG, DY, EM, HTK

Project administration: AMR, ALW, DW, MJD

Supervision: DW, MJD

Writing – original draft: TPWD

Writing – review & editing: All authors

## Data Availability

Raw read data have been deposited in the European Nucleotide Archive (ENA), accession number ERP160386. Per-chromosome Zarr files containing SNP and CNV data, annotations and accessibility masks will be available from the MalariaGEN Vector Observatory (https://malariagen.github.io/vector-data/) upon publication. Code required to reproduce the bioinformatic workflows and analysis in this study is available at: https://github.com/tristanpwdennis/AsGARD.

## Supplementary Materials for

### Materials and Methods

#### Sample collection, processing, and sequencing

##### Sample collection and storage

Per-sample collection methods, dates, and GPS coordinates are detailed in **Table S1**. **Afghanistan:** Mosquito larvae were collected between 2014 and 2017(*66*), and raised to adults. Exact per-sample dates are not available.

**Djibouti:** Adult mosquitoes were collected using a combination of BG-Pro and CDC-light traps, at five sites around Djibouti City, in August 2023. In the metadata, the GPS coordinate is the centroid for Djibouti City.

**Ethiopia:** Larval mosquitoes were collected in 2023 using dippers and pipettes from aquatic habitat in plastic jars, subsequently transported to temporary field insectaries and transferred into enamel trays and provided with larval food to raise to adulthood. The trays were checked each day for pupation and pupae were collected into beakers containing water and placed in rearing cages with cotton balls soaked in a 10% sugar solution at the top of the cages. All emerging adults were identified morphologically and preserved in an Eppendorf tube with holes in the lid and put in a zip bug with handfuls of silica gel.

**India**: Data are from 15 *An. stephensi* samples from Bengaluru and Mangaluru in southern India sequenced to approx 15X(*67*).

**Iran:** A combination of adult and larval specimens were collected using a combination of pyrethrum spray catches (PSC) (Saravan, Bandar Abbas townships), and larvae raised to adults (Chabahar), between May 2023 and December 2023

**Pakistan**: Adult mosquitoes were collected using pyrethrum spray catches (PSC) in April 2005 in a previous study(*68*).

**Saudi Arabia:** Adult mosquitoes were collected using Black Hole traps from November 2023 to December 2023

**Sudan**: Adult mosquitoes were collected using a combination of PSC, Prokopack aspirators, and Centres for Disease Control and Prevention (CDC) light traps.. The samples were collected from 4,575 houses and their surroundings through 6 rounds of longitudinal entomological surveillance from 61 sites on a monthly basis during the period July 2022 to March 2023 in 9 states in eastern, northern and central zones of Sudan including Khartoum. Further data was collected from 400 households and their surroundings through expanded entomological surveillance from 8 sites in Khartoum, Gezira, Sennar, and White Nile states.

**Yemen**: Adult mosquitoes were collected between February 2021 and February 2023 using CDC light traps.

##### DNA extraction, Quality Control and Sequencing

DNA was extracted from individual whole mosquitoes using the Qiagen DNEasy Blood and Tissue Kit, before being sent to Novogene UK for library preparation and sequencing to a target coverage of 30X using 2×150bp paired reads on an Illumina NovaSeq X instrument.

#### Variant and CNV calling

##### Read mapping and variant calling

Throughout this study, we used the UCI2018 *An. stephensi* assembly reference genome(*69*), downloaded from Vectorbase(*70*). We used the NVIDIA Clara Parabricks Germline pipeline for read mapping and variant calling. The Germline pipeline incorporates BWA-MEM(*71*), GATK’s MarkDuplicates(*72*) and GATK HaplotypeCaller(*73*) to produce individual .g.vcf files, using a NVIDIA GA100 GPU. Individual .g.vcf files were jointly genotyped with GATK’s GenomicsDBImport and GenotypeGVCFs commands across 250 genomic intervals in parallel. The 250 jointly called vcf files were merged into a single, all-sample VCF using GATK GatherVCFs.

##### Variant filtering

In the absence of a ‘truth set’ of variants, we designed hard variant filters using 100,000 randomly selected SNPs from chromosome 2. We were guided by the strategies used by the MalariaGen Vector Observatory(*74*), and GATK hard filtering guidelines(*75*). We used bcftools(*76*) to filter variants to include only:

- SNPs (excluding INDELs or complex variant types).
- Sites with no more than one ALT allele (--max-alleles 2).
- Sites with no more than 10% of sites missing.
- Sites with a Quality Depth (QD) greater than 5.
- Sites with a Mapping Quality (MQ) greater than 40.
- Sites with a Fisher Strand (FS) of less than 20.
- Sites with a Strand Odds Ratio (SOR) of less than 3.
- Sites with a Depth (DP) between 6000 and 13000.

Additionally, we examined per-sample missingness, and produced principal component analysis (PCA) plots of the samples, removing samples with an overall missingness greater than 10%, and samples grouping abnormally in the PCA (often due to high per-sample missingness, or suspected contamination). The total number of variants retained after variant calling and filtering was 8,730,863, 8,714,133, and 1,363,056 variants in chromosomes 2, 3 and X respectively. The final callset was annotated using SNPEff(*77*), with a custom database generated using the FASTA and annotation GFF3 for the UCI 2018 reference genome, downloaded from Vectorbase. The annotated vcf callset of 551 individuals was converted into zarr(*78*) format using sgkit(*79*).

##### Haplotype phasing

We used Beagle v5.5(*80*) to phase the whole callset, with a window size of 10cM, and an overlap of 1cM. In the absence of a genetic map for *An. stephensi*, we assumed a constant recombination rate of 1cM/Mb.

##### Accessibility mask

We constructed an accessibility map for the UCI 2018 reference genome, following the method used for *An. gambiae*(*15*). We selected 100 individuals at random, and computed the following metrics for each position in the reference genome, for each individual:

- Number of reads covering the position (coverage).
- Number of properly paired reads at the position.
- Average read mapping quality at the position.
- Whether the position was masked by the DUST low-complexity and TandemRepeatFinder Tandem Repeat annotations for the UCI 2018 genome.

We classified a reference genome positions as accessible if:

- The position was not masked by DUST or TRF
- Fewer than 0.2% of individuals have ambiguous alignments
- Fewer than 10% of individuals have a mapping quality (mapq) of less than 30
- At most 0.2% of individuals have no coverage
- At most 10% of individuals have at least half or less than the modal coverage
- At most 2% of individuals have at least twice the modal coverage This mask classified 82% of the genome as accessible.

##### Copy Number Variant Calling

We estimated copy number variants (CNVs) according to:(*19*). In brief: we calculated the following statistics in 300bp windows:

- coverage of reads with a mapq of zero
- coverage of reads with a mapq of less than 10
- coverage of reads with a mapping quality greater or equal to 10

We computed the normalised coverage from the read counts, based on expected coverage of each window given its GC content. For each window, we divided the read counts by the mean read count over all autosomal windows with the same GC percentage. When calculating normalising constants, we accounted for potential high copy number variation by excluding windows with extremely high or low coverage, as well as windows with over 10% inaccessible bases, and less than 50% of reads with a mapq of 0.

We then applied a Gaussian HMM to each individual’s normalised windowed coverage data. CNVs were called by locating contiguous runs of at least five windows with amplified copy number state (CNS > 2, or CNS > 1 for the X chromosome in males). Windowed modal CNVs were then interpolated with the annotation set for the UCI2018 genome, to create a table of normalised copy number for every annotated protein coding gene.

#### Population Genetic Analysis

Unless specified otherwise, population genetic data analysis was largely performed using the scikit-allel(*60*), sgkit(*79*), numpy(*81*), pandas(*82*) libraries. Data were plotted using seaborn(*83*), and plotly(*84*), organised in xarray(*85*), stored in in zarr(*78*).

##### Principal component analysis

We performed principal component analysis (PCA) to identify population structure in our samples. We performed PCA on 100,000 randomly selected SNPs, with a minimum minor allele count of 2, from chromosomes 2, 3 and X and, finding no major difference in clustering pattern between chromosomes, retained the PCA from chromosome 2 only. As uneven sample sizes by population can bias PCA clustering signal(*86*), on samples downsampled to contain no more than 30 samples per analysis population and, noting a difference in the positioning of the Sudanese samples (n=231) compared to other populations, presented the downsampled all-population PCA.

##### Estimation of individual admixture proportions

We estimated individual admixture proportions using *HaploNet*(*65*). Haplonet infers fine-scale population structure using dense phased haplotypes. We ran HaploNet on a whole-chromosome phased VCF for chromosome 2, using the default parameters: (https://github.com/Rosemeis/HaploNet).

##### TreeMix & F-statistics

We investigated the relationships between the different populations with TreeMix(*87*). This creates a maximum-likelihood tree based on allele frequency correlations between populations. We used 822000 randomly selected, accessible sites with a minor ac > 1 from chromosome 2. A block size of 100k was used to account for linkage disequilibrium, and the 1,000 bootstraps were run by resampling blocks of 100 SNPs. The resulting trees were summarized with the sumtrees function in ape(*88*). In the absence of an obvious outgroup, we did not root the tree.

##### Neighbour joining trees

We inferred neighbour-joining trees were inferred 100,000 randomly selected SNPs from chromosomes 2, 3 and X. City-block distances were inferred between all pairs of sites in all individuals. The resulting distance matrix was used to infer a neighbour-joining tree in the anjl(*89*) library, which implements the neighbour-joining algorithm of Saitou and Nei(*90*).

##### Local PCA

Population structure may vary across the genome(*63*). In the case of mosquitoes, heterogeneity may arise as a consequence of chromosomal inversions, which can be detected by performing PCA in windows across the genome(*28*, *63*). We performed local PCA in windows of 50kb across chromosomes 2, 3 and X.

##### Mitochondrial phylogeny

We extracted whole mitochondrial genome consensus sequences for all samples in the dataset by using *bcftools consensus.* We aligned sequences using MUSCLE(*91*), and manually inspected and trimmed them to remove majority-gap regions in AliView(*92*). We inferred a maximum-likelihood phylogeny using IQ-TREE(*64*), automatically selecting a nucleotide substitution model using MODELFINDER(*93*), evaluating support with 1000 ultrafast bootstrap replicates(*94*). For the cytochrome-c-oxidase (*COI*) tree, we performed the same steps, trimming the alignment to contain only the region of the *COI* gene. Trees were plotted with ape(*88*) in R.

##### Fst

We calculated average genomewide Fst between all chromosomes, using allele counts from all population pairs, in scikit-allel using Hudson’s Fst estimator, chosen because it is more robust to unequal sample sizes(*95*).

##### Thetas

We calculated nucleotide diversity (π) and Tajima’s D, in 10kb windows across chromosome 2, using scikit-allel, which, when given the chromosome start and end, calculates π using all sites (including uncalled sites) as a denominator(*96*).

##### ROH

We inferred per-individual and per-chromosome runs of homozygosity (ROH) using scikit-allel, which uses a hidden markov model (hmm) to infer ROH. Output was combined and aggregated to provide per-individual genome-wide estimates of the number of ROH in an individual genome, and the fraction of the genome in ROH. We filtered ROH to include only segments longer than 100kb(*15*, *97*), to include only those putatively a result of recent inbreeding.

##### Mitochondrial phylogenetic tree

We reran the variant calling pipeline above on the *An. stephensi* mitochondrial genome, setting the ploidy to 1, for HaplotypeCaller. The resulting set of SNPs was filtered according to the same set of filters as above. Consensus sequences for each individual were extracted using bcftools. We also included *An. stephensi* mitochondrial genome sequences used in previous publications(*11*). (OR607949.1, OR607950.1, ON421572.1, ON421574.1, OK663480.1, OM801691.1, OM801703.1, OM801693.1, OM801697.1, MH651000, OK663481, OK663479, OK663482, OQ865406.1, OQ865407.1, MW197100.1, MW197099.1, MW197101.1, OM865140.1, PP387838.1, MF975729.1, MF975728.1, MF124611.1, MF124610.1, MF124609.1, MF124608.1, MG970565.1, MK170098.1, KF406694.1, KF406693.1, KF406701.1, KF406680.1, KX467337.1, MH538704.1, MK726121.1, MN329060.1, LR736015.1, LR736014.1, LR736013.1), as well as the *An. maculatus* mitochondrial genome sequence (KT899888.1). We created a multiple sequence alignment (MSA) with MUSCLE(*91*), and manually inspected and edited the alignment in AliView(*92*). We used the resulting MSA as to infer a maximum-likelihood phylogeny with IQTREE 2(*64*), using in-built ModelFinder(*93*) to automatically estimate a nucleotide substitution model (HKY+F+I), evaluating support with 1000 ultrafast bootstrap replicates(*94*). We chose *An. maculatus* as an outgroup taxon for rooting, as it was the closest taxonomic relative to *An. stephensi* with a published mitochondrial genome. For the analyses of the *COI* sequences, we downloaded all of the sequences referenced in(*48*), and aligned them to the MSA. We manually trimmed the alignment to include only the core *COI* sequence, and further trimmed gap-containing sequences, to a final alignment length of 318bp. We estimated a phylogeny as per the previous whole mitochondrial genome tree.

#### Spatial population genetic analysis and modelling

##### Spatial population structure

The distribution of genetic variation across space provides an insight into dispersal processes, and therefore into invasion dynamics. We sought to infer landscape genetic structure in *An. stephensi* using fEEMS(*62*), a software library for inferring ‘migration surfaces’. 100,000 SNPs were randomly selected from chromosomes 2 and 3, and used as input for fEEMS. Triangular grids, of spacing 75km, spanning the analysis region (Horn of Africa and the Arabian Peninsula) were created using the ddgridR(*98*) library. The resulting grid, dataframe of sampling locations, and PLINK format SNPs, were used as input for fEEMS. Leave-one-out cross-validation was performed as per the fEEMS user manual, and the grid with the lowest cross-validation error was selected for plotting. The final weighted grid object was exported and plotted in R.

Based on the PCA, admixture proportion barplot, TreeMix, and fEEMS results, we hypothesised that the sampled *An. stephensi* population in the Horn of Africa comprises two separate invasion events - one in Ethiopia and Djibouti, and another in Sudan. We further explored this hypothesis by analysing the relationship between linearised Fst and geographic distance (inferred using distHaversine in geosphere) between sampling locations, between nucleotide diversity (π) and geographic distance (inferred using distHaversine in geosphere(*99*)) to the primary container port for Sudan (Port Sudan) and Ethiopia (Port of Djibouti), using generalised linear models. Full details of model summaries, formula, and distributions, are given in **Table S3**. Various families of distribution were explored, and goodness-of-fit established by comparing Akaike Information Criteria and residual diagnostics, using the DHARMa(*100*) library.

##### Environmental drivers of An. stephensi dispersal

Geographical and environmental features shape how organisms move across landscapes. Understanding how landscape connectivity may contribute to *An. stephensi* dispersal and life-cycle has been the subject of much interest, as this understanding can be used to model future spread of *An. stephens*i(*4*, *31*, *48*). We explored the impact of environmental heterogeneity on *An. stephensi* genetic connectivity by using a landscape genetics approach to identify environmental variables that influence gene flow. Landscape genetics studies tend to model the effect of environmental distance and genetic distance, between pairs of sampled locations(*56*). In this study, we used linearised Fst as the genetic distance matrix, between sites in Djibouti, Ethiopia, and Sudan. To generate environmental distances between sampling sites, we used Circuitscape(*101*), an approach that borrows from electrical circuit theory to predict connectivity in heterogeneous landscapes(*33*). Although more computationally expensive than least-cost-path and straight-line distances, circuit theory considers all possible paths connecting populations across a landscape, and has greater explanatory power for predicting gene flow(*33*).

##### Environmental data - sources and rationale

We downloaded gridded raster data for 10 different environmental variables spanning the invasive range of *An. stephensi* in Sudan, Ethiopia and Djibouti covered by this study (a bounding box of (xmin = 25, xmax = 85, ymin = 2, ymax = 37). **Table S4** details the data links, time-spans, units, and resolutions. All raster data were scaled to values between 0 and 1 prior to analysis.

- **Relative humidity (hurs)**. Rainfall poorly predicts *An. stephensi* abundance, instead being shaped by patterns of temperature and land use(*31*). Instead, the role of relative humidity on epidemic malaria in Indian urban areas suggests that this parameter may influence landscape permissivity for *An. stephensi*(*32*). We downloaded relative humidity data spanning 1981-2011 from CHELSA (climatologies at high resolution for the earth’s land surface area)(*102*).
- **Mosquito development rate (mdr).** Bionomic studies of *An. stephensi* suggest an optimal temperature window for development(*103*). Temperature data can therefore be used to derive a distribution of mosquito development rate (mdr). We downloaded raster data for average annual mean temperature from CHELSA, spanning 1981-2011, and applied a transformation based on the distribution of mdr as a function of temperature.
- **Human transit friction (friction**). In *An. stephensi*, human transit hubs have been hypothesised as more genetically connected(*11*). We downloaded the Land-based travel speed friction surface from the Malaria Atlas(*104*). The friction map enumerates land-based travel speed, expressed as minutes required to travel one metre, per pixel, for the nominal year 2015(*105*).
- **Human population density (pop).** *An. stephensi* is an urban malaria vector. The presence of high densities of humans, and urbanisation, may therefore influence landscape connectivity. We downloaded data for human population density from WorldPop(*106*).
- **Enhanced vegetation index (evi)**. Vegetation cover may provide an outdoor resting place for mosquitoes, serve as fodder for livestock - also hosts of *An. stephensi* - and indicate areas of higher human habitation in the landscape. We downloaded data on enhanced vegetation index - a measure of greenness derived from satellite reflectance data from the Moderate Resolution Index Spectrometer (MODIS).
- **Elevation (ele)** and **slope** (**slope**). Organism movement may be hindered by mountainous and rugged terrain. We downloaded data on elevation, as a measure of metres above sea level, and slope, as a measure of the average angle of terrain in a grid cell, from the Terra Advanced Spaceborne Thermal Emission and Reflection Radiometer (ASTER) Global Digital Elevation Model (GDEM)(*107*).
- **Cattle density (**cattle). *An. stephensi* is zoophilic, and exhibit a strong preference for livestock as a source of blood(*108*). We downloaded data on cattle density from the United Nations Food and Agriculture Organisation (FAO) Gridded Livestock of the World (GLW) database(*109*), 2020 version.
- **Irrigation** (irrigation). *An. stephensi* breeds in irrigation related habitats, such as irrigation canals, pits, dams and channels(*110*). We downloaded gridded irrigation data from the UN FAO(*111*), where each pixel enumerates the percentage area equipped for irrigation around the year 2005.
- **Mean wind travel time** (mean.wind.traveltime). Wind facilitates long and short-range mosquito dispersal(*112*). The direction and extent of this influence is determined by wind speed and direction. We used the windscape R package(*113*), to determine the mean wind travel time between sampling locations in the HoA. Windscape converts hourly wind rasters into landscape connectivity graphs, then uses the graphs to estimate directional wind flow among sites of interest. Windscape enables the use of high-frequency time series wind rasters, as opposed to a snapshot of wind conditions, enabling capture of temporal variability in wind conditions. We downloaded wind data from the European Centre for Medium-Range Weather Forecasts Reanalysis, version 5 (ERA5)(*114*). ERA5 covers a 31km grid resolution, and combines climate observations from meteorological stations and satellites to produce detailed estimates of past climate. We downloaded data for wind *u* and *v* at 10m, spanning the study area, for the 1,5,10,15,20 and 25th of each month between 2012-2023, between the period spanning dusk in the Horn of Africa (17:00, 18:00, 19:00 and 20:00), when *An. stephensi* is the most active. We chose ERA5 over other sources of wind data (e.g. the climate forecast system reanalysis - CFRS) due to its temporal resolution spanning the period of *An. stephensi* invasion, as well as higher spatial resolution.

##### Environmental data - modelling

Our modelling approach followed the established framework for identifying landscape variables influencing genetic distance, sometimes called *Isolation-by-resistance* or *Isolation-by-environment*. These approaches take measures of genetic distance, typically Fst, and use Mantel tests of paired distance matrices, or linear regression, to establish relationships between genetic distance and various environmental covariates(*56*) In this analysis, we used linearised Fst(*115*) as a genetic distance measure. We calculated the straight-line distance, least-cost path distance (using the R package *gdistance*(*116*)), and resistance (using the Circuitscape package), between all pairs of points within Sudan, Ethiopia, and Djibouti. After initial examination, the resistances produced by Circuitscape, a method that treats a landscape like an electrical circuit, and calculates environmental distance as ‘resistance’ across all possible dispersal paths(*33*, *101*), produced the clearest relationships between genetic distance, and all environmental covariates. Circuitscape can determine multiple different routes for dispersal whereas straight-line or least-cost path distances do not capture this potential variability so we reasoned that Circuitscape likely produced the most biologically plausible results for this analysis. For wind, we calculated mean wind travel time between points using Windscape (see above). Our genetic vs environmental resistance data consisted of single values giving the linearised Hudson’s Fst between sampling locations with more than 5 samples, and environmental distance (resistance in the case of covariates analysed with Circuitscape, and mean travel time with wind) for each pair of points. We subset our data into two invasion populations, based on our hypothesis, supported by the genetic analyses, that the Horn of Africa *An. stephensi* samples in this study constitute two geographically distinct populations – one for Sudan, and one for Ethiopia and Djibouti. We only included linearised Fst estimates from within each hypothetical invasion population, omitting between-population comparisons. Thus, like our Fst values, the explanatory variables give a single value between two locations allowing us to explore their predictive potential. Examining the output of Circuitscape revealed that the irrigation and cattle analyses produced many NA values due to missing data in the rasters, and these were subsequently dropped from the analysis. Finally, we calculated the geographic distance between points, to explore whether the environmental and landscape covariates are correlated with genetic distance after controlling for distance limited dispersal (isolation-by-distance).

We started by parameterising two Bayesian Gaussian linear regression models using *rstanarm*(*117*), to select priors. We compared a Student’s *t* prior (a reasonable default prior when effects have a chance of being large), and a regularised horseshoe prior (RHS) (enforces strong shrinkage for small effects, ideal for collinear and highly dimensional data)(*118*). The models contained all predictors (**Table S5**). The RHS prior produced a model with greater predictive accuracy using leave-one-out-cross validation (LOO-CV)(*119*).

We then performed predictive variable selection(*120*) using the *projpred* library. Briefly, the variable selection process splits the model into submodels, and performs LOO-CV on each submodel, ranking the predictive performance of each covariate. The predictive performance of the variables is shown in **Figure S8**. This process showed that the variables **mdr, evi** and **friction** contributed to explaining variability in the dependent variable Fst, while additional covariates added little, allowing us to select these 3 explanatory variables as covariates to include in the final model.

We re-ran the regression using the same parameters and RHS prior as above, including the three selected predictors. We tested whether this model was a better fit than a model of isolation-by-distance alone, by running a regression of Fst and pairwise distance. Furthermore, we determined whether the predictors varied significantly by country, by adding an interaction term for “invasion”, corresponding to the two spatially segregated populations of Ethiopia & Djibouti, or Sudan (**Table S5)**. Model comparison using LOO-CV indicated that a model including three environmental covariates as predictors of Fst was preferred over models that included only pairwise distance or models that incorporated interactions between environmental covariates and invasion status. (**Table S5**).

#### Resistance gene characterisation

##### Resistance allele frequencies

We downloaded coordinates for candidate IR genes of interest (*Vgsc, Rdl, Ace1, Cyp9k1*), based on their involvement in resistance in other *Anopheles* vectors of malaria(*121*), from VectorBase. We extracted SNPs spanning these coordinates (**Table S2**) from the annotation database (see ***functional annotation***), calculated and plotted their frequencies per population.

##### Selection

We performed genome-wide selection scans (GWSS) for evidence of recent selection by using Garud’s H-statistics. Because of small per-population sample sizes in India, we omitted these populations from the analysis(*122*). GWSS data were calculated in windows of 1000 haplotypes, aggregated and plotted by population.

We estimated selection and gene flow at highlighted IR loci in the genome by using hierarchical clustering of haplotypes. This approach is used to detect gene flow and selection at loci of interest(*15*). We selected haplotypes spanning the *Vgsc* (3:42817709-42817709), *Rdl* (3:8345440-8348441), *Ace1* (2:60916071-60917000) and *Cyp9k1* (X:9721225-9722225) genes, as well as those spanning the *Cyp6* (2:67473117-67501071), *Gste* (3:70572788-70584603) and *Coeae* (3:18784698-18801820) gene clusters. (**Table S2**). We chose these regions based on the presence of selective sweeps in the GWSS, as well as their role in insecticide resistance in other *Anopheles* taxa. We used a hierarchical clustering approach implemented using *scipy*(*123*) to cluster haplotypes.

##### CNV analysis

We selected genes based on a) evidence of selection in the H12 scan and haplotype clustering analyses, b) evidence of involvement in resistance in other *Anopheles* taxa, and c) prior evidence of copy number variation in other *Anopheles taxa.* We selected genes from the *Cyp6, Gste, Coeae, CoaeJH* clusters, and *Ace1* and *Cyp9k1* genes, and extracted syntenic orthologs between *An. stephensi* and *An. gambiae* in VectorBase, based on the start and end of regions containing genes annotated as members of the same gene family (e.g. confirmed or putative cytochrome p450s in regions containing *Cyp6* orthlogs in the *An. gambiae* PEST genome on VectorBase). This resulted in a table of 29 genes (**Table S2**). We plotted the modal copy number for these genes as a heatmap in R.

### Supplementary Text

#### Insecticide resistance architecture of invasive An. stephensi

Despite widespread resistance to all widely used insecticide classes in invasive *An. stephensi*, the genomic architecture of resistance is very poorly understood. We performed a comprehensive survey of loci implicated in resistance in *Anopheles* taxa. We screened our data for missense mutations in the target-site resistance genes: *Vgsc*, *Ace1* and *Rdl* (**Figure S7, Table S2**):

- **Ace1** confers resistance to organophosphates and carbamate insecticides. G94S, a recently discovered *Ace1* mutation in *An. stephensi*, which is not currently known to be associated with a resistance phenotype(*44*), is present at frequencies of 0.25-0.97 in the HoA, Yemen, and India, but at 0.06 Afghanistan and Pakistan, and is absent from KSA and Iran. (**Figure S7**).
- ***Rdl*** A296S, conferring resistance to organochlorine insecticides(*13*) is almost fixed in KSA (0.98-1), while present at low to intermediate frequencies in all other populations (0.07-0.55) (**Figure S7)**.
- The ***Vgsc*** knock-down-resistance (*kdr*) mutation L1012F/S, commonly known as L1014F/S due to its amino acid codon in *Musca domestica*, where it was first discovered(*124*), confers resistance to DDT and pyrethroid insecticides. *Kdr* L1012S is present only in Afghanistan and Pakistan(*42*), at a frequency of 0.12. L1012F was present at low to intermediate frequencies in the HoA and Yemen (0.07-0.5), and KSA (0.11-0.7). Both L1012F and S are present at low frequencies (0.08-0.06) in Iran, and are absent from India.

To identify other regions of the genome potentially involved in insecticide resistance, we performed per-population genome-wide selection scans (GWSS) using Garud’s H12 statistic, omitting India (INM and INB) due to small (<10) cohort sizes. (**Figure 2a**). Signals of H12 in invasive populations in the Horn of Africa, Yemen, and KSA-Riyadh, are very noisy genomewide compared to their native range counterparts in Afghanistan/Pakistan and Yemen, presumably due to reductions in genome-wide diversity as a consequence of recent introduction. This is particularly marked in Sudan, Yemen and KSA-Riyadh. There are signals of selective sweeps at IR-associated loci in all populations, but with marked differences among them (**Table S2, Figure 2a**). In KSA Eastern (SAE), H12 is elevated in regions containing *Rdl,* the *Cyp6* cytochrome p450 cluster, and the *Gste* cluster. In KSA Riyadh (SAR), there are signals of sweeps at *Rdl,* and *Gste* (**Figure 2a, Table S2**). In Iran, there are signals of sweeps at *Gste* (IRH) and *Cyp6* (IRS). In Afghanistan and Pakistan, signatures of a selective sweep are present at the *Cyp6,* carboxylesterase alpha-esterase *(Coeae),* carboxylesterase JH *(CoeJH),* and *Gste* clusters, as well as at the *Cyp9k1* gene **(Table S2, Figure 2A**). Signals of sweeps at *Gste* and *Coeae* are also present in India, and apparent sweeps at the *Cyp6, Coeae* and *Gste* clusters are apparent in populations from the Horn of Africa and Yemen, though less clear in Sudan and Yemen due to noisy background signals. *Ace1* appears to be under selection in the HoA and Yemen, but not elsewhere. Sweep signals are absent in all populations at the voltage-gated sodium channel (*Vgsc*) gene.

To further explore patterns of selection at loci under selection in one or more populations we present the results of hierarchical haplotype clustering (**Figure S5),** aligned with the sample population. We analysed the *Cyp6, Gste* and *Coeae* clusters, as well as the *Rdl, Cyp9k1* and *Ace1* genes, based on their involvement with IR on other mosquito taxa and evidence of selection in one or more *An. stephensi* populations (**Figure 2a**). We include the *Vgsc* gene due to its pivotal role in resistance in other mosquito taxa, despite absence of a selective signal (**Figure 2A**), or association with resistance in previously published data(*37*). We found evidence of strong selective sweeps (multiple haplotypes with low or zero distance from one another) in all genes except for *Cyp9k1* and *Vgsc* (**Figure S5**).

Sweeps in *Ace1* and the *Cyp6, Gste* and *Coeae* loci are shared between invasive *An. stephensi* populations and, in all cases, sweep clusters contain mixes of haplotypes from the Horn of Africa and Yemen, and Afghanistan and Pakistan, indicative of gene flow of these haplotypes between these populations (**Figure 4b-e**). In all cases, Indian and KSA populations form distinct clusters (**Figure 1b-e, Figure S5**), and evidence of selection at the *Rdl* locus is apparent in KSA populations (**Figure 2a**). We do not observe striking signatures of selective sweeps at the *Vgsc* locus in any population (**Figure 2a**) (note that samples from Sudan form clades of identical haplotypes, due to substantially reduced haplotype diversity in this population). These data suggest that pyrethroid resistance in *An. stephensi* in the Horn of Africa and Yemen may be conferred predominantly through *Ace1* and metabolic mechanisms, as opposed to other target-site mechanisms such as *Vgsc* and *Rdl*, and that selected haplotypes originate in Afghanistan and Pakistan, consistent with this being the closest potential origin population.

In *An. gambiae* and *An. funestus*, copy number variants (CNVs), lead to elevated expression levels, and sometimes also changes in protein sequence, and are widely associated with resistance, particularly at metabolic genes. There is some evidence of CNVs associated with detoxification genes in *An. stephensi*(*13, 58, 125*), but to date, no systematic, genomewide analysis of CNVs in samples across the native and invasive range has been performed. Given evidence of strong selection at *Ace1* and various metabolic loci in the Horn of Africa and Yemen, we performed a genome-wide search for copy number variation (**Figure 2b**,). We searched the results for three gene families: *Ace1, Coeae, CoeJH, Cyp6, Cyp9k1 and Gste* (**Table S2**), based on evidence of selection in the HoA, Yemen, Afghanistan and Pakistan. We find evidence of extensive copy number variation in the *Coeae, Cyp6 and Gste* clusters (**Figure 2b**), with modal copy number reaching 10 in the *Gste* and *Coeae* clusters in the HoA and Yemen. KSA predominantly contained CNVs in the *Gste* cluster, with a small number in *Coeae*. Afghanistan and Pakistan, the HoA and Yemen contained very similar CNV profiles at *Coeae*, *Cyp6* and *Gste*. India contains CNVs at the *Coeae*, *Gste* clusters, and *Cyp9k1* (**Figure 2b**). Similarly to the selection scans and clustering dendrograms, KSA and India are the most different, whereas Afghanistan and Pakistan, the HoA, and Yemen, share similar CNV profiles.

## Supplementary Figures

**Figure S1:**
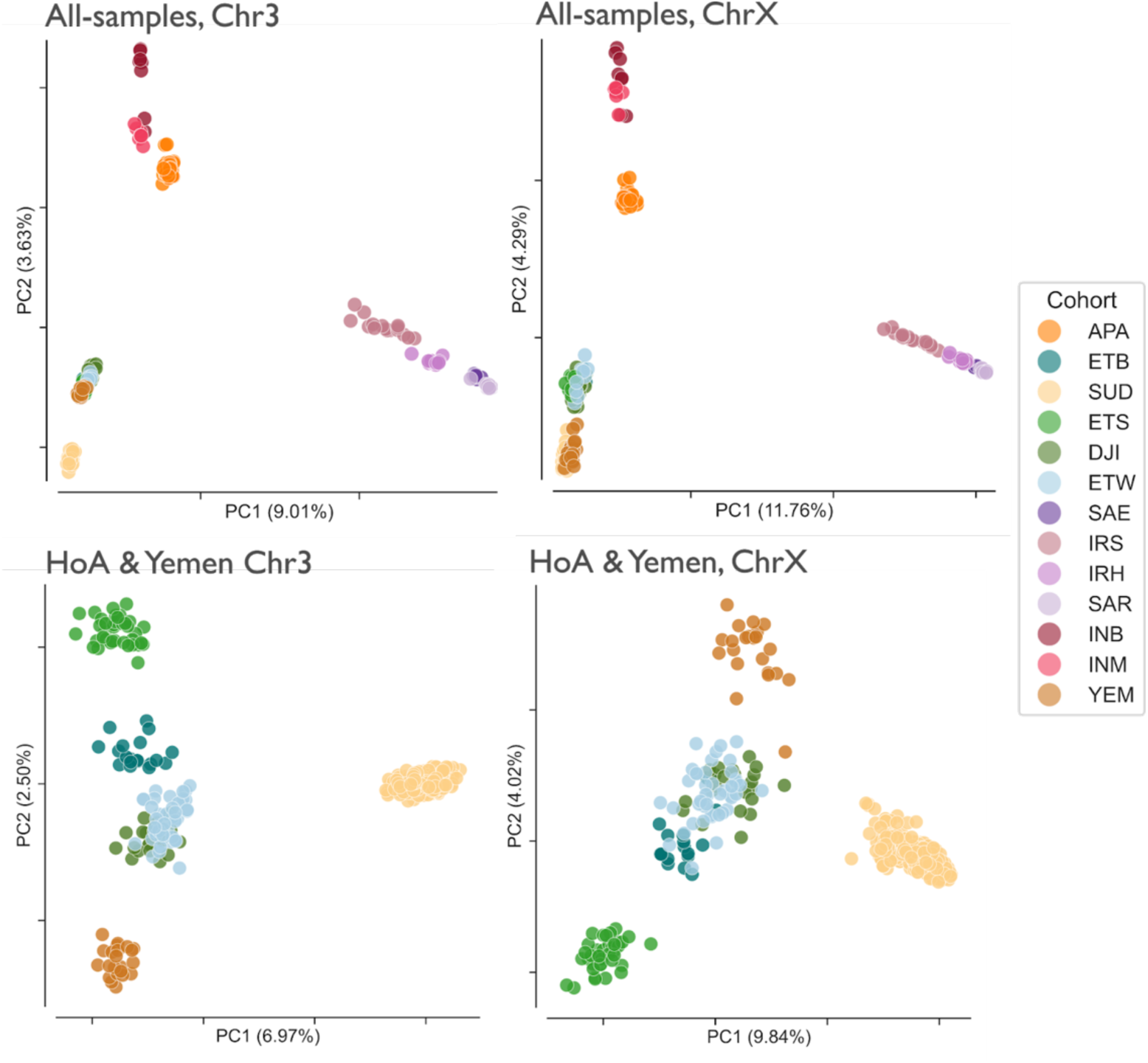
Principal component analysis of chromosomes 3 and X from all populations (top row), and HoA & Yemen populations (bottom row). Percentage of total variance explained by each PC shown in brackets. PCAs A and B were downsampled to no more than 30 samples per cohort, while C and D used all individuals in all HoA and Yemen cohorts (see Supporting Material). PCAs were computing with 1e5 randomly chosen accessible SNPs from their respective chromosomes. Cohort indicated by point colour (see key).

**Figure S2:**
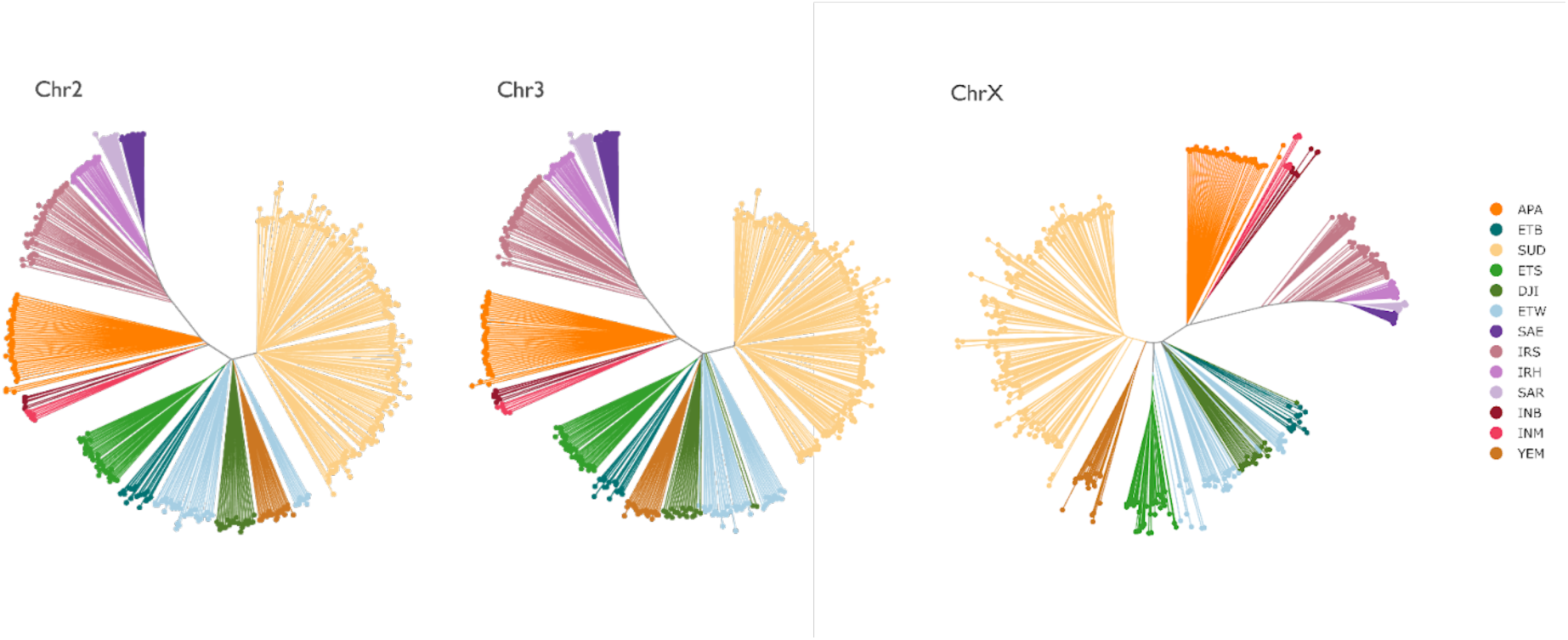
Neighbour-joining trees of *An. stephensi* chromosomes 2,3 and X. Branch length indicates city-block distance between samples (tips). Tip points coloured by population (see key). Trees were inferred using 1e5 randomly chosen accessible SNPs from each chromosome.

**Figure S3:**
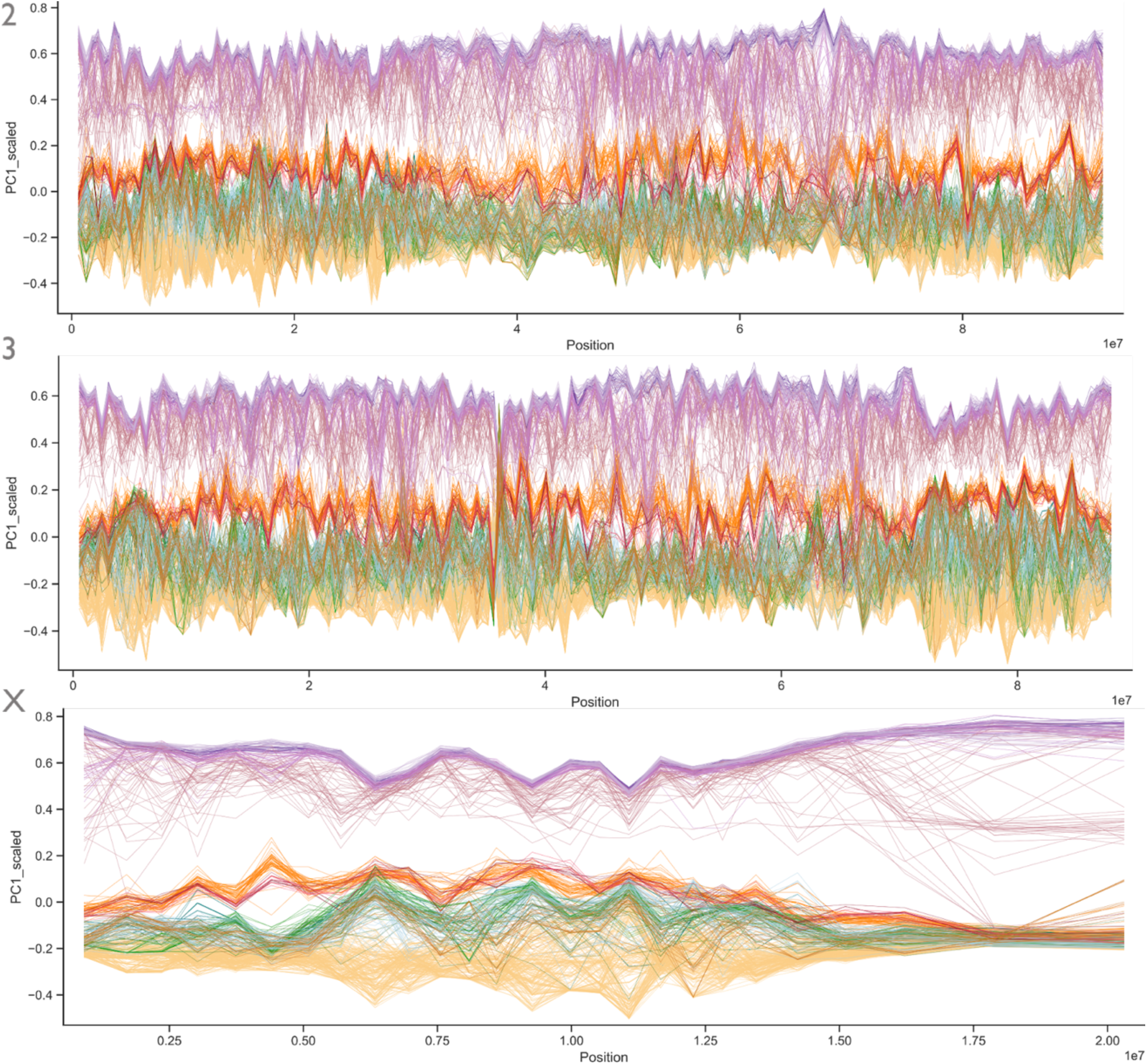
No inversion polymorphisms detected in *An. stephensi*. PC1 (y axis) of principal component analysis (PCAs) calculated using accessible SNPs in 50kb windows for all samples across chromosomes 2, 3 and X, with individual lines indicating sample, coloured by population (see key). In the presence of an inversion polymorphism, PCA will usually reflect the karyotype (e.g. homozygous negative, heterozygous, homozygous positive), as opposed to the neutral structure of the population(*28*, *63*). The plots above show an absence of structuring by inversion karyotype.

**Figure S4:**
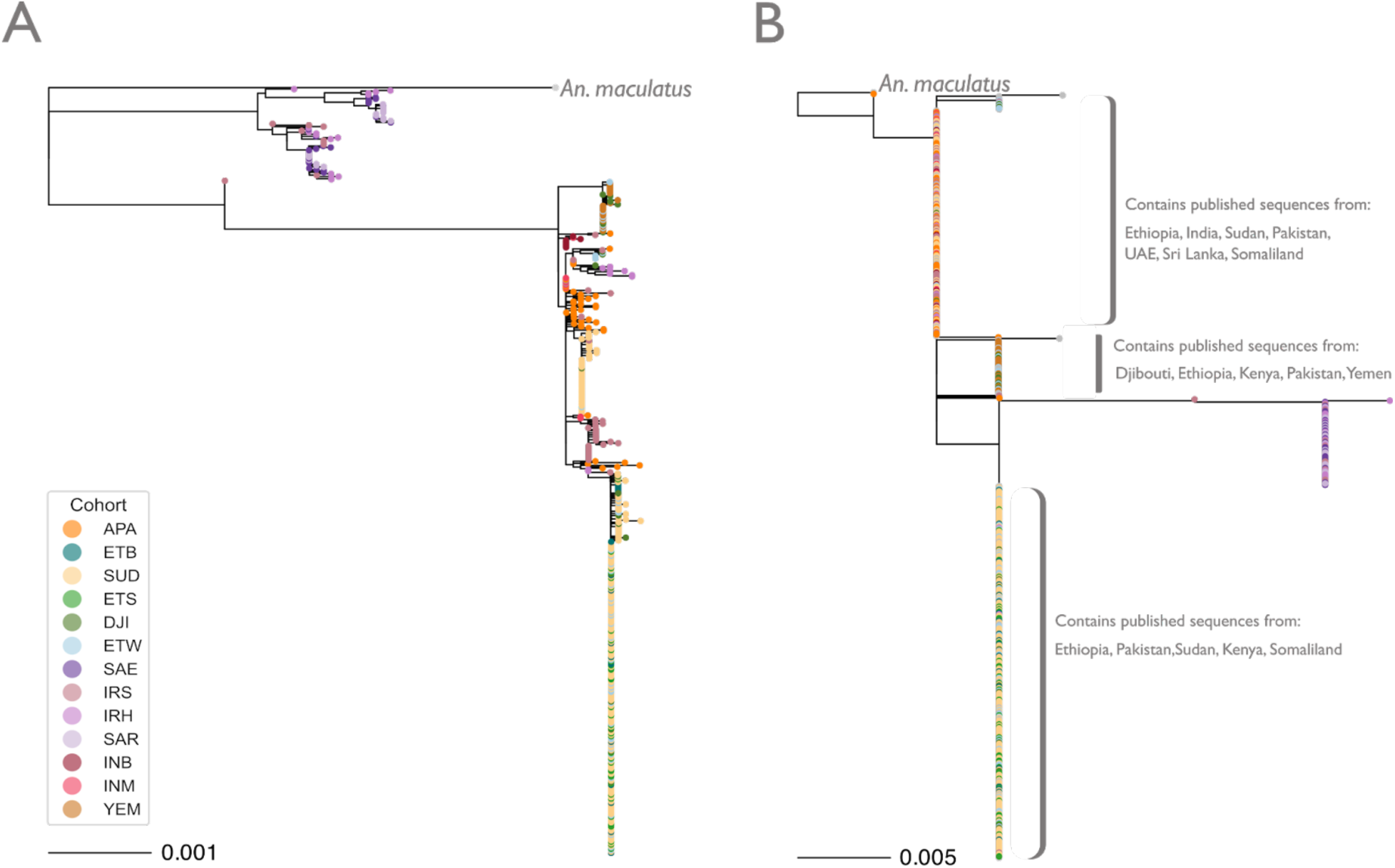
Mitochondrial (A) and *COI* (B) phylogenies. (A) was inferred from whole mitochondrial genome alignments of all samples in this study. (B) was inferred by subsetting the whole-mitochondrial alignment in (A) to the region containing the *cytochrome-b-oxidase (coi)* gene, then realigning with published *An. stephensi* COI sequences(*48*) (Methods). Grey tips in (B) indicate published sequences, with text detailing origin countries of published sequences in each clade. Coloured tips are samples from this study, coloured by cohort (see key). Scale bars indicate substitutions per site. Trees were inferred using maximum-likelihood, with automated model selection (HKY+F+I) for both. Support for each tree was evaluated with 1000 ultrafast bootstrap replicates in IQTREE(*64*). Trees were rooted on the *An. maculatus* mitochondrion (A) and *coi* (B). The outgroup branch was artificially shortened for ease of plotting.

**Figure S5:**
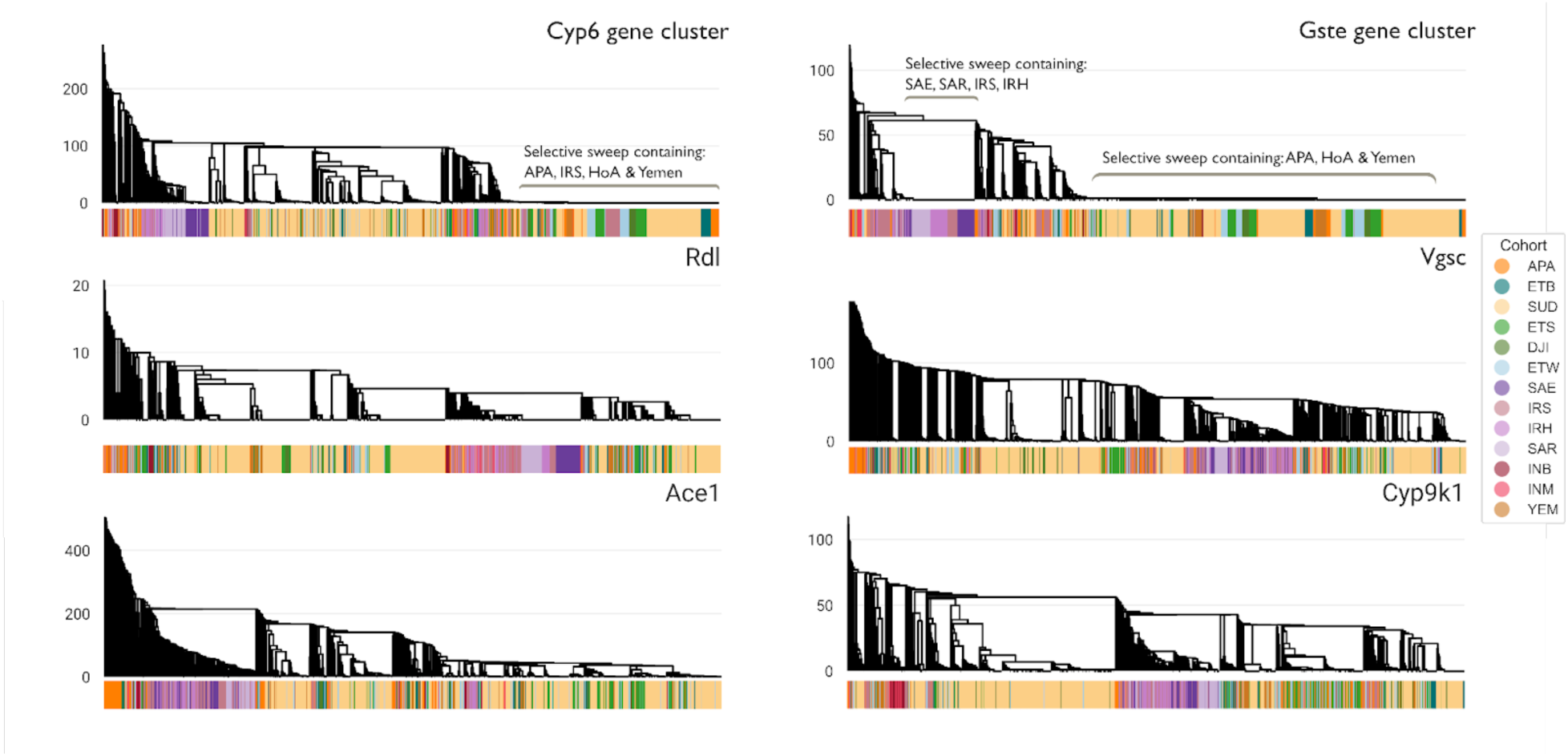
Patterns of gene flow at swept resistance haplotypes. Haplotype clustering dendrograms of insecticide resistance gene families. Gene cluster (*Cyp6* or *Gste*) and gene name (*Rdl, Vgsc, Ace1, Cyp9k1*). Indicated in top right of the panel. Y axis (branch length) indicates distance in number of haplotypes between adjacent tips. X axis (tips) indicates individual haplotypes, with the bar below the plot matching the cohort from which that haplotype was sampled. Legend to the right indicates cohort. Specific sweeps are annotated with text and brackets. Dendrograms were created by computing a distance matrix between phased haplotype counts between all individuals. Haplotypes were taken from the start and end of the first and last gene of a gene cluster, for *Cyp6* and *Gste* clusters, and from the start and end of the region containing the longest transcript of a single gene for *Rdl, Vgsc, Ace1* and *Cyp9k1*.

**Figure S6:**
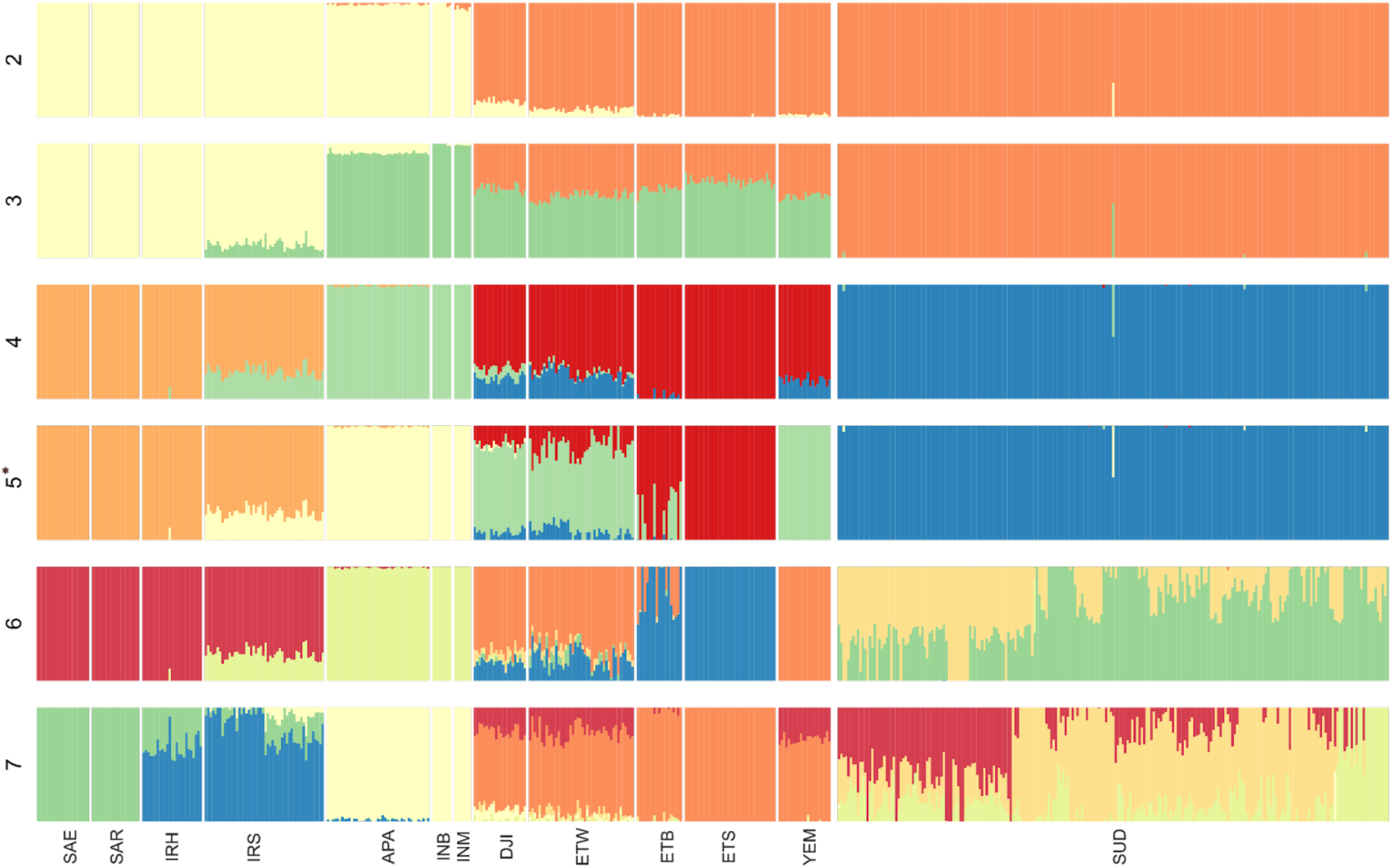
Individual admixture proportions for all cohorts. Barplots indicate the respective contribution of *K* notional ancestral populations (see row label), per individual (stacked bar colour). Barplots are panelled row-wise by *K*, column-wise by cohort. Admixture proportions were inferred with *HaploNet*(*65*) on phased haplotypes across the entirety of chromosome 2.

**Figure S7:**
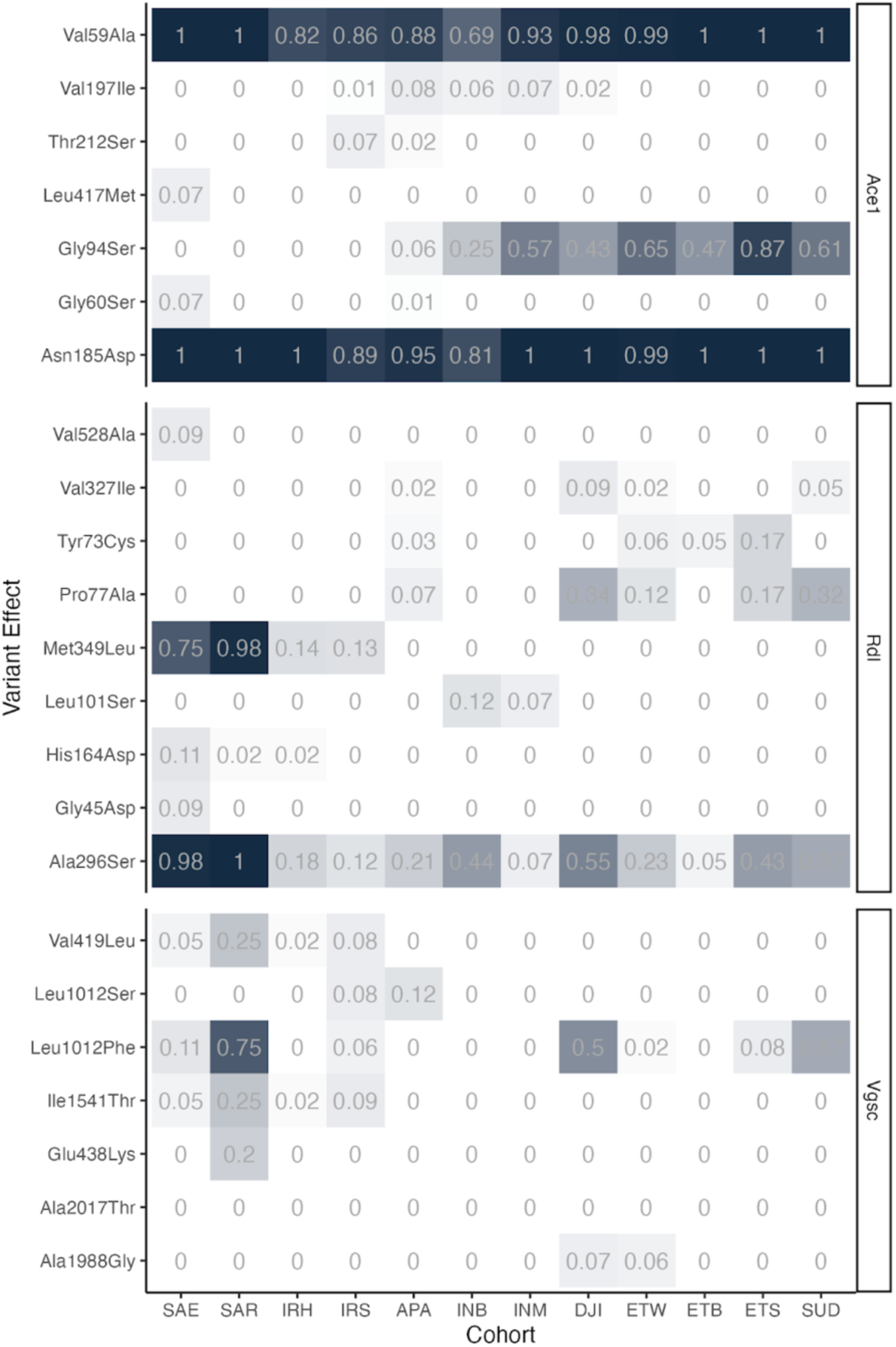
Allele frequencies in target-site resistance genes, *Ace1, Rdl* and *Vgsc*. Y axis denotes variant effect: amino acid position and change, X axis denotes cohort. Plot is panelled row-wise by gene. Frequencies were computed from accessible non-synonymous SNPs with a frequency over 0.01 in at least one cohort.

**Figure S8:**
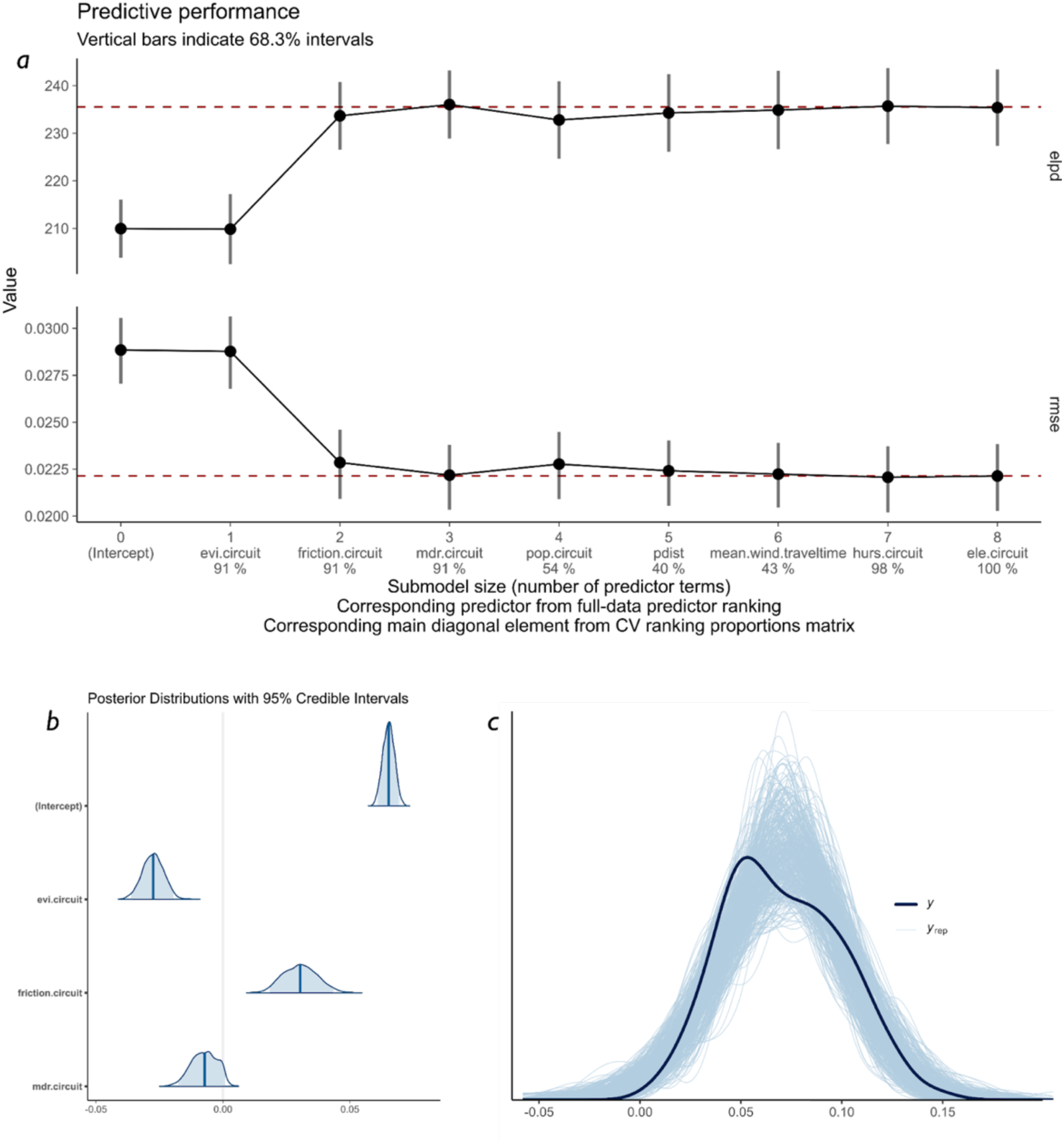
Statistical model selection to associate environmental variables with genetic dispersal (using linearised Fst as a proxy), model coefficient distributions and visualisation of model fit to data. **(a)** Variable selection for the final model fitted to selected environmental covariates to predict linearized Fst (**Table SX**). Predictive variables included are scored to indicate the degree of contribution that each makes to explaining the variability in the Fst value. Selections are either made using the expected log predictive density (elpd) [top panel] or root mean square error (rmse) [lower panel], and evaluated using a leave-one-out cross-validation (LOO-CV) approach. Once variables reach the indicated dashed red lines, variables are no longer informative. Thus, both approaches suggest wind travel time, environmental vegetation cover, and surface friction are key explanatory variables. **b** shows the posterior distributions for coefficients of predictors for the final selected submodel (**Table S5**), with the mean and 95% credible intervals shaded in dark and light blue, respectively (y axis). **c** shows the result of a posterior predictive check showing the kernel density estimate (kde) for the data (dark blue line) and 1000 model draws (light blue lines).

